# A spatial multi-scale fluorescence microscopy toolbox discloses entry checkpoints of SARS-CoV-2 variants in Vero E6 cells

**DOI:** 10.1101/2021.03.31.437907

**Authors:** Barbara Storti, Paola Quaranta, Cristina Di Primio, Nicola Clementi, Nicasio Mancini, Elena Criscuolo, Pietro Giorgio Spezia, Vittoria Carnicelli, Giulia Lottini, Emanuele Paolini, Giulia Freer, Michele Lai, Mario Costa, Fabio Beltram, Alberto Diaspro, Mauro Pistello, Riccardo Zucchi, Paolo Bianchini, Giovanni Signore, Ranieri Bizzarri

## Abstract

We exploited a multi-scale microscopy imaging toolbox to address some major issues related to SARS-CoV-2 interactions with host cells. Our approach harnesses both conventional and super-resolution fluorescence microscopy and easily matches the spatial scale of single-virus/cell checkpoints. We deployed this toolbox to characterize subtle issues related to the entry phase of SARS-CoV-2 variants in Vero E6 cells. Our results suggest that in these cells the variant of concern B.1.1.7, (aka Alpha variant), became the predominant circulating variant in several countries by a clear transmission advantage. In fact, in these cells B.1.1.7 outcompetes its ancestor B.1.177 in terms of a much faster kinetics of entry. Given the cell-entry scenario dominated by the endosomal “late pathway”, the faster internalization of B.1.1.7 could be directly related to the N501Y mutation in the S protein, which is known to strengthen the binding of Spike receptor binding domain with ACE2. Remarkably, we also directly observed the main role of clathrin as mediator of late-entry endocytosis, reconciling it with the membrane localization of the ACE2 receptor previously attributed to caveolin-enriched rafts. Overall, we believe that our fluorescence microscopy-based approach represents a fertile strategy to investigate the molecular features of SARS-CoV-2 interactions with cells.

## INTRODUCTION

SARS-CoV-2 has rapidly spread worldwide generating a pandemic with devastating social consequences. The development of a handful of novel and effective vaccines^1^ represented a brilliant scientific achievement and it holds promise for a rapid end of the pandemic. Nonetheless, the way out of pandemic could be slowed by the emergence of novel SARS-CoV-2 lineages endowed with better ability to spread and infect humans while featuring lower *in vitro* susceptibility to neutralizing monoclonal and serum antibodies^2^. In this context, elucidation of structure-property relationships that modulate virus-cell host checkpoints, such as entry, replication, and egress, is crucial to assess the role of genome mutation on virus infectivity.

SARS-CoV-2 contains four structural proteins, namely spike (S), envelope (E), membrane (M), and nucleocapsid (N) proteins (Scheme 1a). S is a ∼180 kDa glycoprotein anchored in the viral membrane and protruding as homotrimers from the viral surface (the “corona”)^3^. S plays the most important roles in viral attachment, fusion and entry^4,5^. The N-terminal S1 subunit contains the receptor binding domain (RBD) that mediates SARS-CoV-2 binding to the cell membrane receptor ACE2^6^. The C-terminal S2 subunit (Scheme 1b) is responsible for the fusion of the viral envelope with cellular membranes to deliver the viral RNA^7^. The S-mediated membrane fusion follows two proteolytic events: i) the “priming” cleavage that occurs at the S1/S2 interface, which yields S1 and S2 non-covalently bound in a pre-fusion conformation, and ii) the “activation” cleavage that occurs within the S2 subunit (S2’) to trigger the fusion process^8^. For several CoVs, including SARS-CoV-2, fusion can occur either at the plasma or endosomal membrane, according to the “early” and the “late” pathways of entry^9^. Availability of the transmembrane-bound protease TMPRSS2 favors virus entry through the early pathway^10^. Conversely, in TMPRSS2-negative cell lines, CoVs internalize by the late pathway and fusion is triggered by the Cathepsin B/L proteases^9^. Quite remarkably, SARS-CoV-2 bears a polybasic amino acid insert (PRRA) at the S1/S2 junction (Scheme 1b). This site is potentially cleavable by furin, a protease commonly found in the secretory pathway of most cell lines^11^. Accordingly, a few studies suggest that SARS-CoV-2 may be primed at S1/S2 by furin (or related proteases) during maturation, thereby harboring a cleaved S protein during egress and secondary infection^12, 13^. The cleaved S protein seems to be advantageous for activating the early pathway in TMPRSS2-expressing cells^14^.

**Scheme 1.**
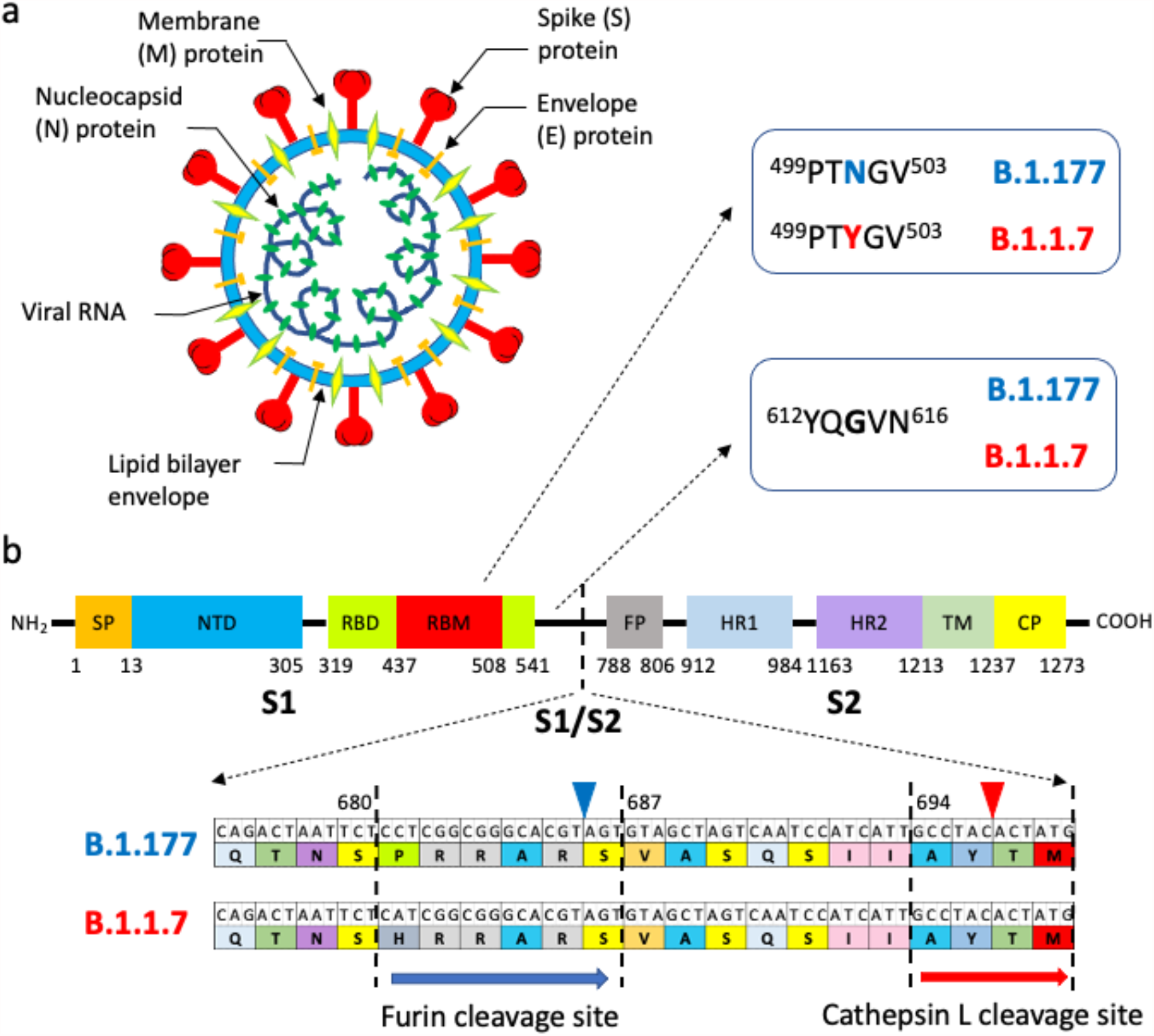
Structure of SARS-CoV-2 and differences between B.1.177 and B.1.1.7 in the Spike protein. (a) SARS-CoV-2 structure. (b) Scheme of S protein. S is composed of the S1 and S2 subunits, which are further subdivided into SP: short peptide, NTD: N-terminal domain, RBD: receptor binding domain, RBM: receptor Binding Motif, FP: fusion peptide, HR1-2: repetitive heptapeptides, TM: transmembrane domain, CP: cytoplasmic peptide. Beside the common D614G mutation, the two most relevant mutations of B.1.1.7 with respect to B.1.177 are reported: N501Y in RBM and P681H at the S1/S2 cleavage site (indicated by the dashed line). On bottom, genome (GenBank accession number NC_045512 used as reference genome) and primary sequence map of the S1/S2 boundary for virus isolates belonging to B.1.1.177 and B.1.1.7 lineages. The proteolytic cleavage site 1 (furin) and 2 (Cathepsin L) are enclosed into dashed lines ^15^, and the relevant proteolyzed linkages are highlighted by blue and red wedges, respectively.

The mechanistic knowledge of virus entry in cells is relevant for developing drugs tailored to prevent infection^16 17^. For instance, therapeutic strategies aimed at inhibiting TMPRSS2 protease activity are currently under evaluation^18, 19^. The proposed, yet unclear and non-exclusive involvement of clathrin and caveolin-1 as mediators of endocytosis in the late pathway may afford further molecular targets to stop pathogenesis^20^. Nonetheless, mutations in the S protein may be crucial for the first step of viral transmission of novel SARS-CoV-2 variants, with significant epidemiological consequences. In particular, D614G became a dominant mutation in SARS-CoV-2 lineages that have been circulating worldwide since spring 2020. Indeed, mutation D614G seems associated with a selective infectivity advantage^21^, possibly due to a more favorable entry phase^22^ compared to wild-type SARS-CoV-2. A further evolutionary step of the virus occurred in the late 2020 in the UK, where the B.1.1.7 SARS-CoV-2 variant (recently denominated as “Alpha” variant by WHO) was firstly detected^21^. B.1.1.7 rapidly outcompeted older D614G strains in many countries^23 24^, due to its 40-70% higher transmissibility that could be associated with ∼30% higher mortality rates^25^. Although B.1.1.7 retains D614G, the enhanced transmissibility of this lineage appears to be related to two additional mutations in the S protein: 1) N501Y, which resides in the Receptor Binding Motif of S, and 2) P681H, which is next to the furin cleavage S1/S2 site (Scheme 1b). Yet, the exact mechanism that confers dominance to B.1.1.7 over the older D614G strains is still unclear, although it has been proposed that N501Y may increase the affinity for ACE2 receptor^26, 27^ and/or P681H may modulate the amount of cleaved S protein harbored by infecting viruses thereby influencing their entry mechanism^28, 29^.

Recent advances in fluorescence microscopy opened the way to individual virus imaging as a tool to understand viral life cycle. The dynamic and heterogeneous nature of virus-cell interactions is the perfect framework for highly sensitive imaging systems such as confocal fluorescence microscopy and Total Internal Reflection Fluorescence (TIRF) microscopy. Of note, TIRF enables imaging of a 100-150 nm layer above the coverslip where 2D cell cultures are adhered, it is thus ideally tailored to follow dynamic processes occurring at the cell membrane like viral entry. Yet, viruses such as CoVs have a size around 100 nm, i.e. well below the optical resolution of confocal and TIRF microscope on the focal plane (200-300 nm), and details of single viral particles interacting with subcellular structures may be only partially revealed with these techniques. Optical super-resolution methods that break the light-diffraction barrier either by leveraging on the photophysical properties of the fluorescent probe or by structuring the excitation light, may easily reach the 20-150 nm spatial scale^30^. Indeed, STimulated Emission Depletion (STED) and Single Molecule Localization Microscopy (SMLM) have been recently applied to image single viruses of different families at <100 nm, also in the cellular context^31, 32, 33^. To our knowledge, however, no super-resolution imaging of full (or pseudotyped) SARS-CoV-2 interacting with cells was yet described in the literature.

In this study, we deploy for the first time a multi-scale fluorescence microscopy toolbox to investigate entry checkpoints of SARS-CoV-2 with two general goals: 1) demonstrate that imaging SARS-CoV-2 at single virus level does help answering biological questions that can only be partially addressed by *in vitro* techniques, and 2) highlight the ability of super-resolution techniques to afford morphology details of virus structure and molecular interactions with the cell. Our multi-scale toolbox was organized according to the resolution capability of each technique: confocal and TIRF microscopy (200-300 nm) were applied to visualize interactions at cell level; super-resolution microscopy techniques (structured illumination in airyscan mode: 120-180 nm, STED: 70-100 nm, SMLM: 25-40 nm) were applied to reveal single-virus morphology and interactions with cell substructures. By our approach we shed light on the different entry kinetics of variant B.1.1.7 compared to B.1.177, an older D614G lineage with large diffusion in Europe in late 2020,^34^ as well as on the role of clathrin and caveolin in mediating the first endocytic step in the late pathway in Vero E6 cells. Beside their own relevance, we believe that our results are representative of a new and fertile approach for the study of SARS-CoV-2 interactions with cells.

## RESULTS

### 1. Virus isolation and setup of imaging toolbox

Two different clinical isolates of SARS-CoV-2, B.1.177 and B.1.1.7, were used for all experiments. The B.1.177 (hCoV-19/Italy/LOM-UniSR10/2021, GISAID Accession ID: EPI_ISL_2544194) and the B.1.1.7 (hCoV-19/Italy/LOM-UniSR7/2021, GISAID Accession ID: EPI_ISL_1924880) strains were isolated on Vero E6 cells from nasopharyngeal swabs from COVID-19 patients at Laboratory of Microbiology and Virology, Vita-Salute San Raffaele University. For safety reasons, SARS-CoV-2 was manipulated in a biohazard safety level 3 (BSL3) and imaging of virus-cell interactions was performed on fixed cells by immunocytochemistry and following indirect labeling. Adherent Vero E6 cells infected by B.1.177 or B.1.1.7 were methanol-fixed and immunostained by anti-S or anti-N rabbit antibodies followed by fluorescently-labeled anti-rabbit secondary antibodies. The use of Alexa488 and Alexa647 dyes was suitable for both confocal/structured illumination (airyscan) and SMLM by the direct STORM approach (dSTORM). Indeed, dSTORM exploits the intrinsic cycling of these fluorophores between bright (on) and dark (off) states to image and localize sparse single molecules at different times across a large field of view and reconstruct a pointillist super-resolved map of the labeled specimen^35^. Conversely, STED nanoscopy requires stable and non-blinking fluorophores because the resolution improvement is performed by the targeted detection of non-depleted fluorophores and high photon flux is necessary^36^. Thus, we selected Atto594 and Atto647 for two-color STED imaging. STED was always performed in the *separation of photons by lifetime tuning* (SPLIT) modality^37^ easily enabled by the Leica Stellaris 8 (Leica Microsystems, Mannheim, Germany) and commercially called τ-STED.

### 2. Kinetics of viral entry

To our knowledge, a detailed comparison of B.1.177 and B.1.1.7 replication kinetics in cell culture is still unreported. Therefore, we infected Vero E6 cells at 0.001 multiplicity of infection (MOI) to enable multicycle replication and the amount of virus in the external medium was checked at different times by reverse transcriptase-polymerase chain reaction (RT-PCR). Vero E6 cells were selected since they represent a common model for SARS-CoV-2 infection^10^. At 24 hours post-infection (hpi), the growth curves of the two viruses were similar; subsequently, however, cells infected by B.1.1.7 were significantly more productive than B.1.177 (Figure 1a).

**Figure 1.**
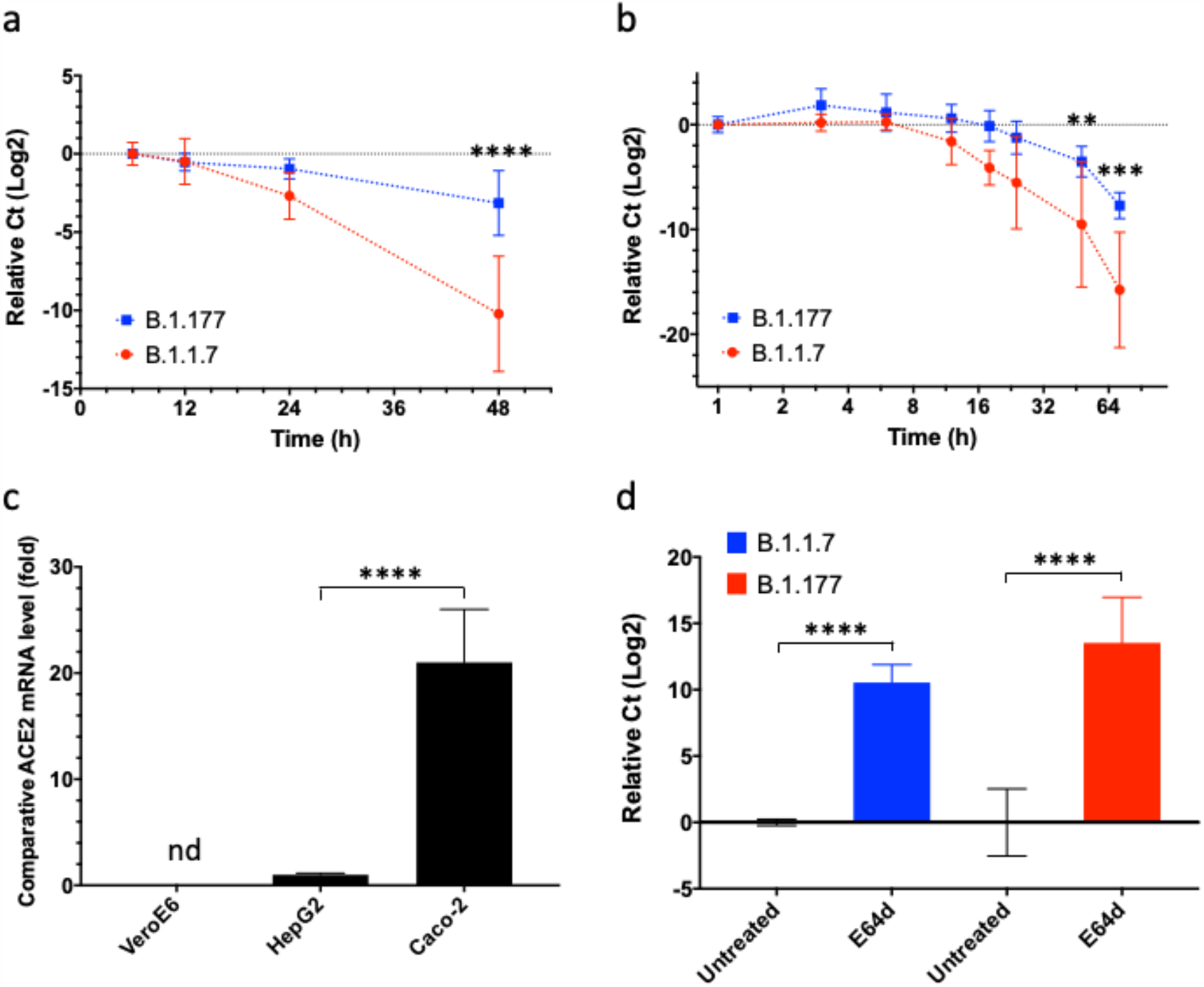
Kinetics of cell entry of B.1.177 and B.1.1.7 SARS-CoV-2. (a) Growth curves showing the release of viral genome into the medium of Vero E6 cells in permanent contact with viruses. (b) Same as (a) but Vero E6 cells were incubated for only 1h with viruses. (c) Expression of TMPRSS2 in total cellular RNA of Vero E6, HepG2, and Caco-2 cells by quantitative real-time PCR. (d) Difference between released viral genome at 48 hpi following treatment with Cathepsin L inhibitor E64d. In (a,b) the viral concentration is expressed by the difference between the cycle threshold (Ct) at the observation time and the average Ct at first time point. In (d) the relative viral load is calculated as the difference between the released genome in untreated and E64d-treated cells.

To better characterize the differences between B.1.1.7 and B.1.177, we modified the infection protocol to avoid the second round of infection by the viral progeny. Cells were incubated with B.1.1.7 and B.1.177 for 1h; then the inoculum was removed, and the cells were washed and maintained at 37 °C for different times (Figure 1b). B.1.1.7 was early detectable in the supernatant and grew with time. Conversely, the external concentration of B.1.177 was negligible up to 24 hpi and became detectable only after 48 hpi. Taken together these data suggest a faster cell entry of B.1.1.7 in tested cells.

To further investigate the entry mechanism of B.1.177 and B.1.1.7 in Vero E6, we checked for the presence of the membrane protease TMPRSS2. In agreement with literature data^38^, we found out that Vero E6 cells express almost no TMPRSS2, whereas control Caco-2 cells do (Figure 1c). In absence of TMPRSS2, SARS-CoV-2 is thought to enter cells by the “late pathway”, i.e. by the endosomal route^13^, whose milestone is the cleavage of S protein by endosomal cathepsins, particularly Cathepsin B/L ^4^. Accordingly, we tested the effect of cathepsin inhibitor E64d on the kinetics of the infection (Figure 1d). At 48 hpi almost no viral production was observable in cells treated with E64d, confirming the key role of cathepsins in virus entry on Vero E6 model and, most importantly, that both B.1.177 and B.1.1.7 do follow the “late pathway” to infect Vero E6 cells.

### 3. Visualization of the virus during entry and egress

To investigate whether the faster infection kinetics of B.1.1.7 was partially attributable to a more rapid entry into cells, we imaged virus-cell interactions at early infection. At 1 hpi with the two viral strains (0.001 MOI), only B.1.1.7 was visible on the periphery of cells (Figure 2, top row). Yet, the fluorescence of B.1.1.7 viral particles almost disappeared from 3h onward (not shown). Conversely, B.1.177 increasingly accumulated on cells with time and at 6 hpi several viral particles accumulated on the cell periphery (Figure 2, bottom row).

**Figure 2.**
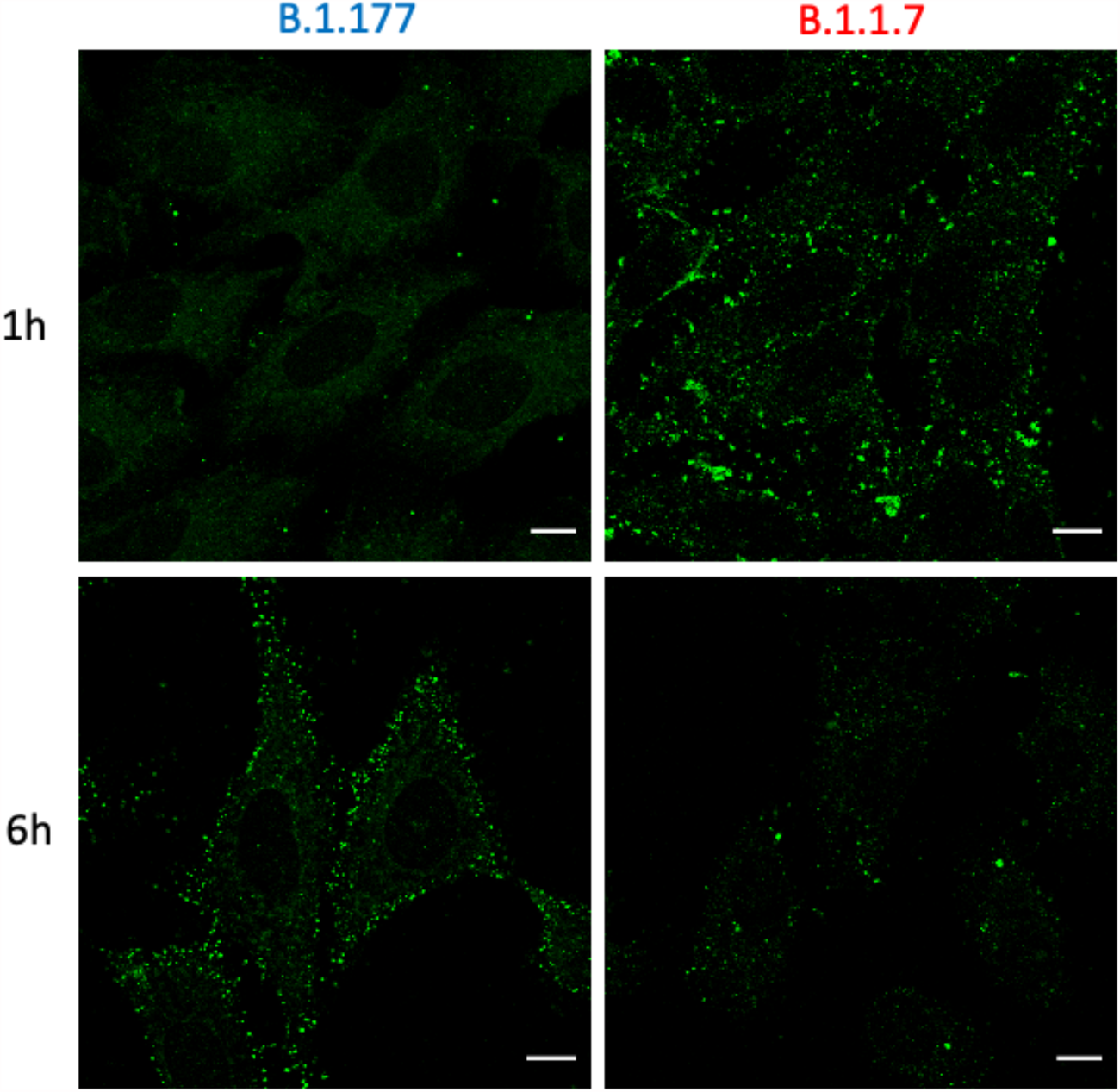
Imaging of cell entry of B.1.177 and B.1.1.7 SARS-CoV-2. Confocal microscopy images of Vero E6 cells at 1 and 6 hpi; Green: S protein, scale bar: 10 µm.

3D imaging suggests that most virions still resided onto or near the plasma membrane (Figure 3). τ-STED microscopy demonstrated that fluorescent particles on the cell membrane were composed of single viruses, with ∼ 100-130 mm size, and very small clusters of about 2-3 virions (Figure 3, inset). These findings are in agreement with a previous electron-microscopy study that showed at 6h membrane-attached single and pairs of wild-type SARS-CoV-2 viruses, with poor cell internalization^39, 40^. The strong interaction of B.1.1.7 with cells at ∼1h detected by microscopy was consistent with the short 1h exposure able to activate strong viral production after one day, at odds with the phenotype observed for B.1.177 (Figure 2b).

**Figure 3.**
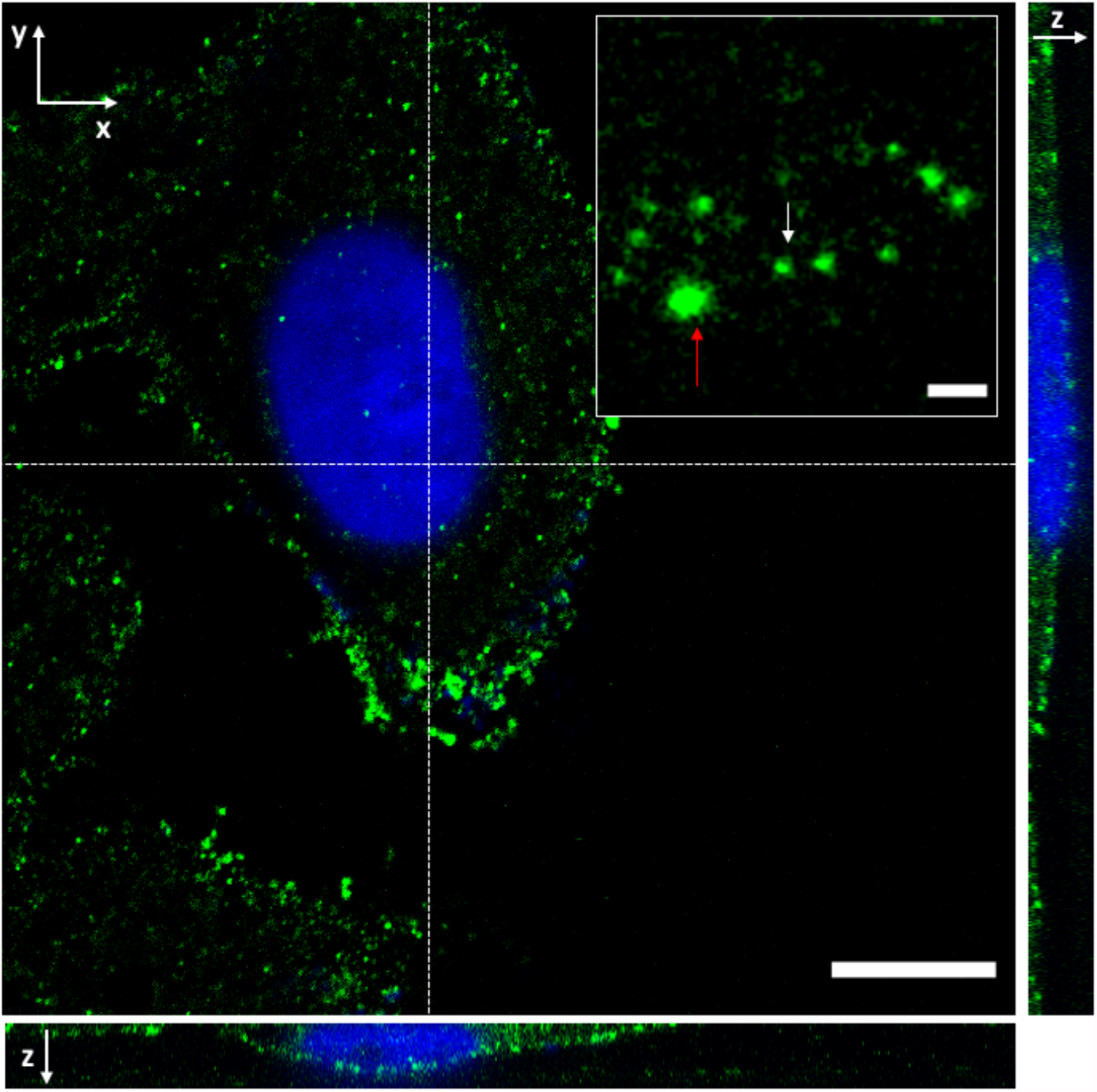
Single B.1.177 virions as well as small aggregates are mostly attached to cell membrane. 3D stack in confocal mode of a Vero E6 at 6 hpi (B.1.177 infection). A medial XY plane of the cell is visible in the large image; on the right and below are reported the YZ and XZ sections corresponding to the dotted line, respectively. Inset: τ-STED imaging revealed that membrane-adhered viral particles consisted of single virions of 100-150 nm size (white arrow) and small viral aggregates (red arrow). Blue: Hoechst 33342, Green: S protein. Scale bar: 10 µm (main panel), 500 nm (inset).

Coherently with a sustained viral replication process, thousands of single virions were discernible in the cytoplasm of Vero E6 at 72 hpi by τ-STED (Figure 4a,c). Large patches of egressing virions accumulated near the plasma membrane, according to the last phase of their viral cycle (Figure 4b). Remarkably, intracellular virions were often found in close proximity and even aligned to microtubules (Figure 4d). This suggests a relevant role for these filaments in the viral cycle, as already demonstrated for other CoVs^41^.

**Figure 4.**
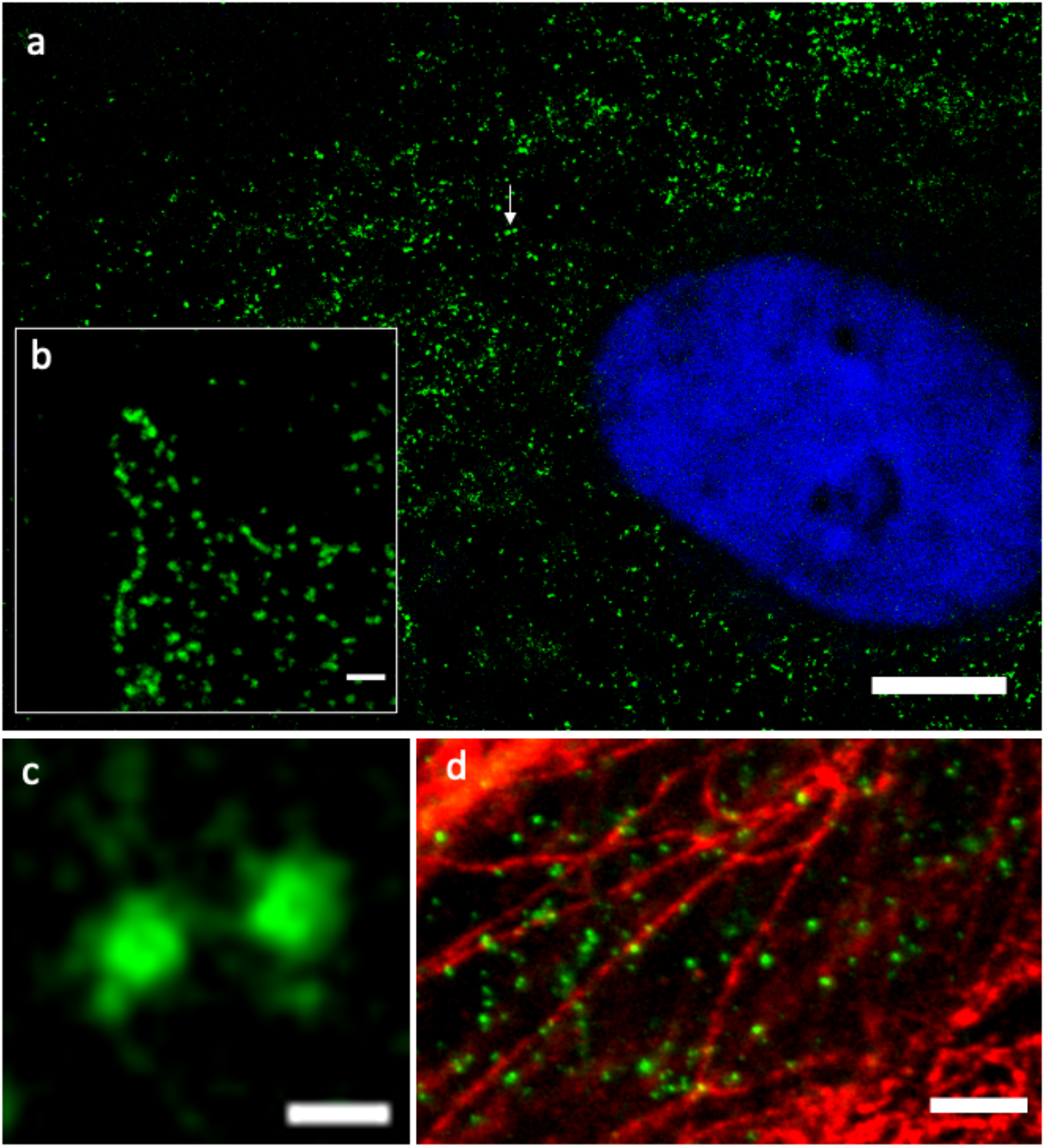
Egressing virions at 72 hpi in a producing cell. (a) τ-STED image of a Vero E6 72 hpi: single B.1.177 virions are diffused throughout the cytoplasm during the egress process. (b) egressing virions ear and along the plasma membrane. (c) Magnified image of two virions whose position is indicated by a white arrow in (a). (d) Single virions near and aligned with microtubules. Blue: Hoechst 33342, green: S protein (a,c,d), N protein (b), red: α-tubulin. Scale bar: 10 µm (a), 500 nm (b), 200 nm (c), 2 µm (d).

### 4. Nanoscale imaging of viral size

Given the nanoscale resolution of dSTORM in the xy plane (average localization precision: 30 nm) and the strong z-sectioning of TIRF imaging mode (100-120 nm), we set out to investigate the size of single virions during their entry (Figure 5a) and egress phases (Figure 5b) by combining dSTORM with TIRF and focusing on the basal membrane plane. To retrieve the viral size, single-molecule localization data were analyzed by density-based spatial clustering of applications with noise (DBSCAN), a clustering algorithm based on localization maps that can discover clusters of arbitrary shapes^42^. Remarkably, in the localization maps DBSCAN identified several clusters (Figure 5c,d) characterized by high labeling density (20,000-600,000 localizations per square μm^2^) that were identified with the single viral particles. Size-distributions of the clusters yielded the average viral radius for both B.1.177 and B.1.1.7 under different labeling conditions (Figure 5e,f). We found out <r> = 50.3±1.6 nm and <r> = 51.4±1.1 nm for B.1.1.7 and B.1.177 when the S protein was labeled, and <r> = 43.9±0.9 nm for B.1.177 when the N protein was labeled. These findings demonstrate that 1) B.1.1.7 and B.1.177 do not differ significantly in size; 2) a smaller virus diameter is detectable when the N protein is labeled, coherently with the location of N proteins inside the virus envelope.

**Figure 5.**
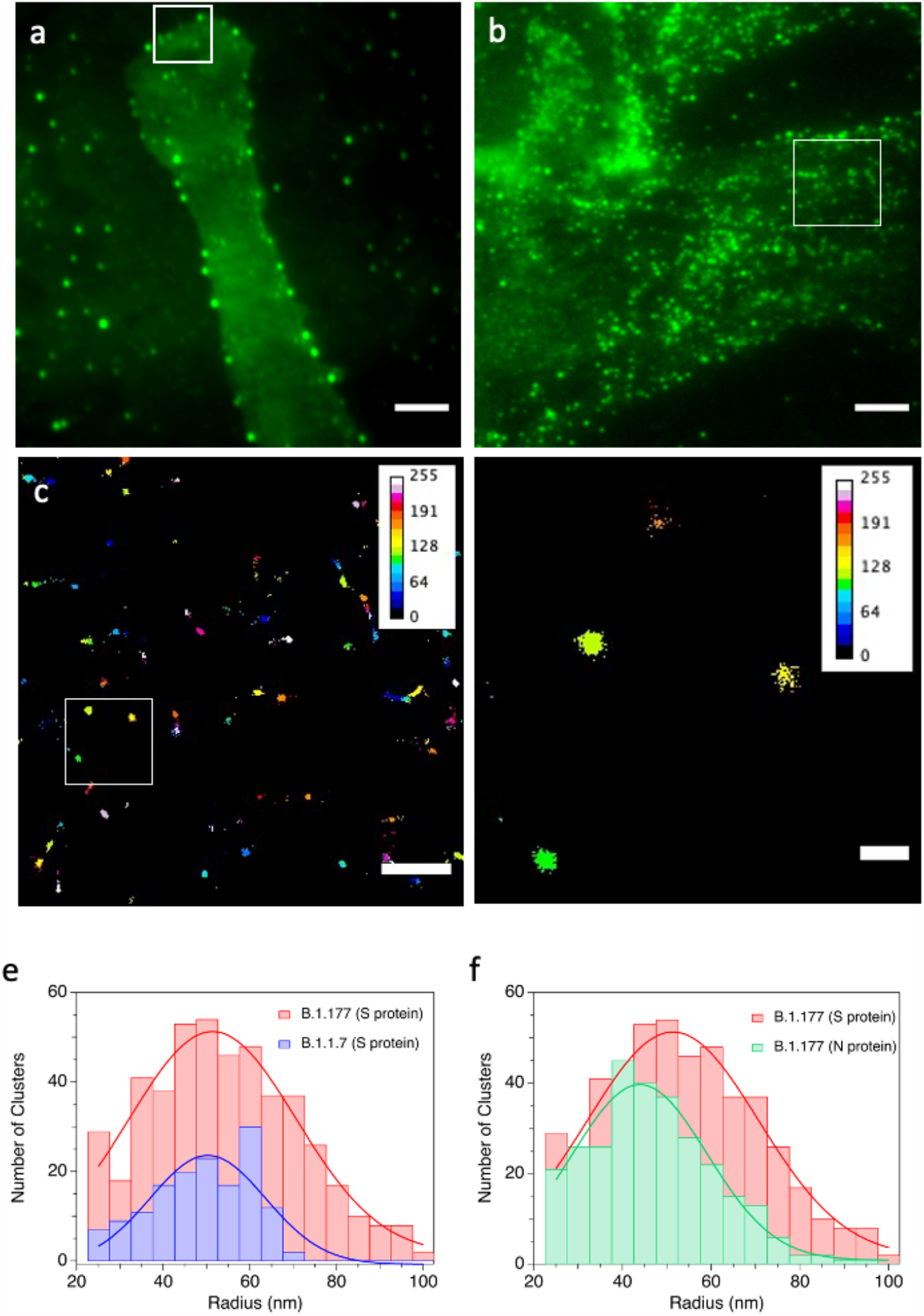
Size of virions by dSTORM-TIRF. (a) TIRF image of a Vero E6 cell at 1 hpi (B.1.1.7 infection).(b) TIRF image of a Vero E6 cell at 72 hpi (B.1.177 infection). (c) Cluster analysis (DBSCAN) of the region of interest (ROI) enclosed in the white squares of (a); clusters are colored according to the relative number of localizations on 0-255 pseudocolor scale; pixel size: 10 nm, average localization precision (±SD): 32±8 nm. (d) Zoom of the region enclosed in the white square of panel (c). (e,f) Distributions of viral particle radius (r) as obtained by cluster analysis: each dataset was fitted by Gaussian curve yielding: <r> = 50.3±1.6 nm, SD: 13.4±2.3 nm (B.1.1.7, S protein), <r> = 51.4±1.1 nm, SD: 19.4±2.6 nm (B.1.177, S protein), <r> = 43.9.4±0.9 nm, SD: 15.2±1.2 nm (B.1.177, N protein). Green: S protein (a), N protein (b). Scale bar: 5 μm (a,b), 1 μm (c), 500 nm (d).

Cryomicroscopy studies have recently highlighted the morphology and dimensions of SARS-CoV-2. Although viruses are not perfectly spherical, the virus envelope has 85-90 nm diameter ^43^. Of note, this figure is in excellent agreement with the size detected by cluster analysis on N-labeled viruses. An average of 26±15 S proteins, about 25 nm long in their perfusion conformation, reside on the surface of each virion, and they can freely rotate around their stalks with an average angle of 40° ± 20° with respect to the normal to the envelope^3^. Thus, the virus “corona” adds an average thickness of about 19 nm to the envelope size, yielding an overall virus diameter of 125-130 nm, i.e. 62.5-65 nm radius.

To check whether our experimental results are in keeping with the published morphology data, we simulated the TIRF excitation of a virion labeled by a primary/secondary antibody couple on its corona and how the localization density of the S proteins resulted on the image plane (Supplementary Information, Figure S1). Given the cylindrical symmetry of the illumination system, we calculated the localization density as a function of the distance (ρ) from the center of a viral particle located at different distances from the basal plane (Supplementary Information, Figure S1), to mimic different experimental conditions. Our simulation showed that the localization density grows up from ρ=0 nm to ρ =48-58 nm, depending on the labeling site on the S protein, to decrease slowly farther off (Supplementary Information, Figure S2). This implies that the maximum fluorescent intensity of an S-labeled virus must be expected slightly above its envelope radius, in good agreement with our cluster analysis results.

### 5. Clathrin-mediated B.1.177 entry mechanism

The early events of the “late pathway” were investigated for B.1.177 by adopting an infection scheme that enabled synchronization of virus entry^44^. Cells were pre-incubated with B.1.177 for 3h at 4 °C, allowing membrane attachment of the virus but preventing its endocytosis. After the chilling step, the non-attached virions were removed, and cells were incubated at 37 °C to promote viral entry. Viral particles were clearly visible near the cell membrane at 2-3 hpi (Figure 6b).

**Figure 6.**
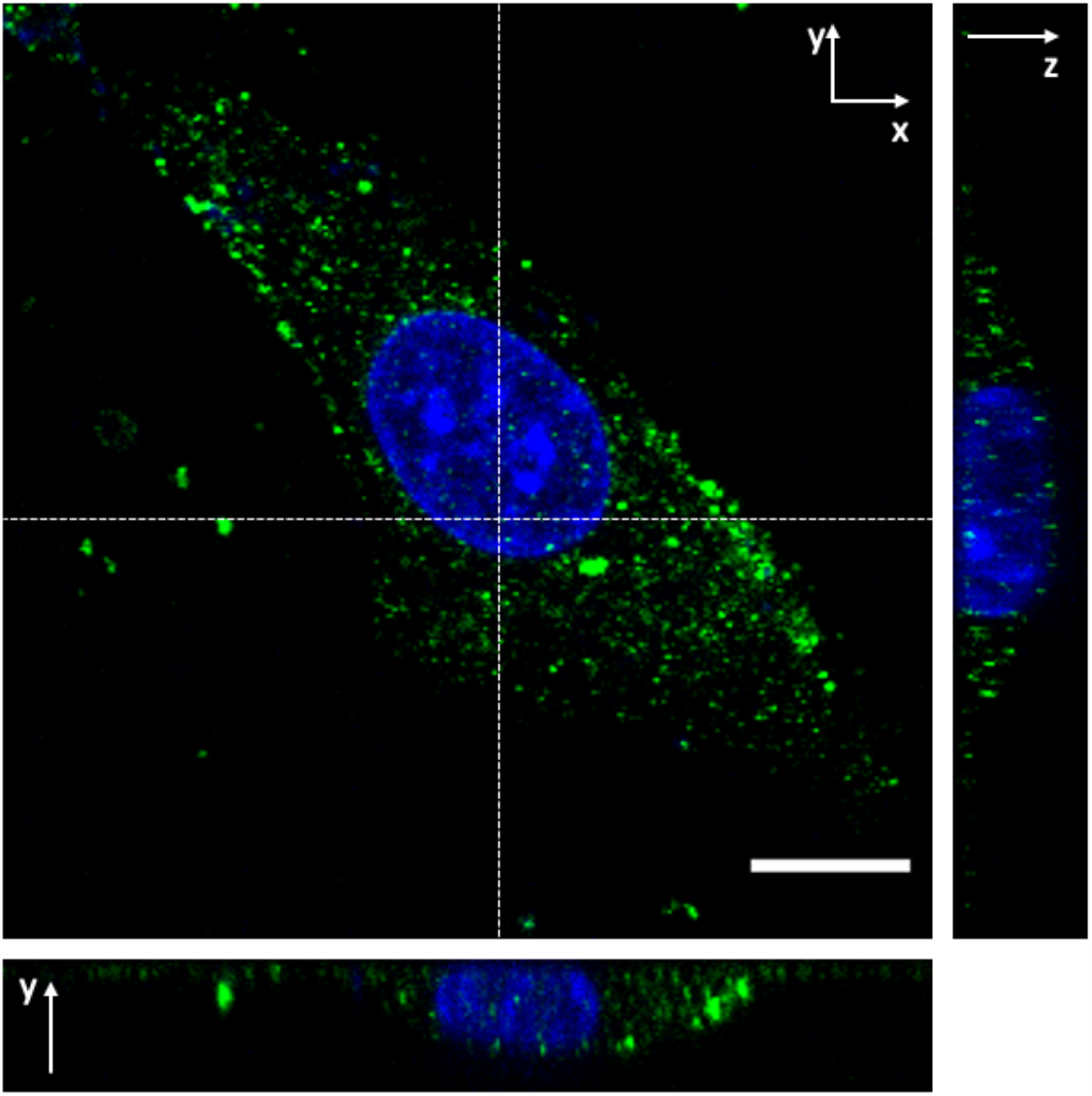
Synchronization of virus entry in Vero E6 cells. Cells were exposed to B.1.177 for 3h at 4 °C, washed and then maintained for 48 h at 37 °C: this procedure led to virus-producing cells, albeit to a lesser extent than control cells, as witnessed by RT-PCR. (c) After 2-3 h post-incubation at 37 °C, most virus are located near the cell plasma membrane, as witnessed by a 3D stack in confocal mode; a medial XY plane of the cell is visible in the main panel; on the right and below are reported the YZ and XZ sections corresponding to the dotted line, respectively; Blue: Hoechst 33342, Green: S protein, scale bar: 10 µm.

Clathrin-mediated and caveolar endocytosis represent the most common initial step of virus endocytosis^45^. Remarkably, dual-color airyscan images alleged a significant colocalization between viral particles and clathrin, but not caveolin-1 (Figure 7). This pattern was quantitatively confirmed by Pearson’s coefficient R, which measures the stoichiometric correlation between the two fluorescent partners as a proxy of their functional association (Table 1).

**Table 1.** Pearson’s coefficients for molecular partners imaged on cell membrane.

|  | ACE2 | RBD | Clathrin | Caveolin-1 | CD71 |
| --- | --- | --- | --- | --- | --- |
| SARS-CoV-2 (S) | - | - | 0.37±0.06 | 0.06±0.04 | - |
| ACE2 | 0.69±0.01 | 0.66±0.07 | - | 0.01±0.05 | 0.15±0.02 |

**Figure 7.**
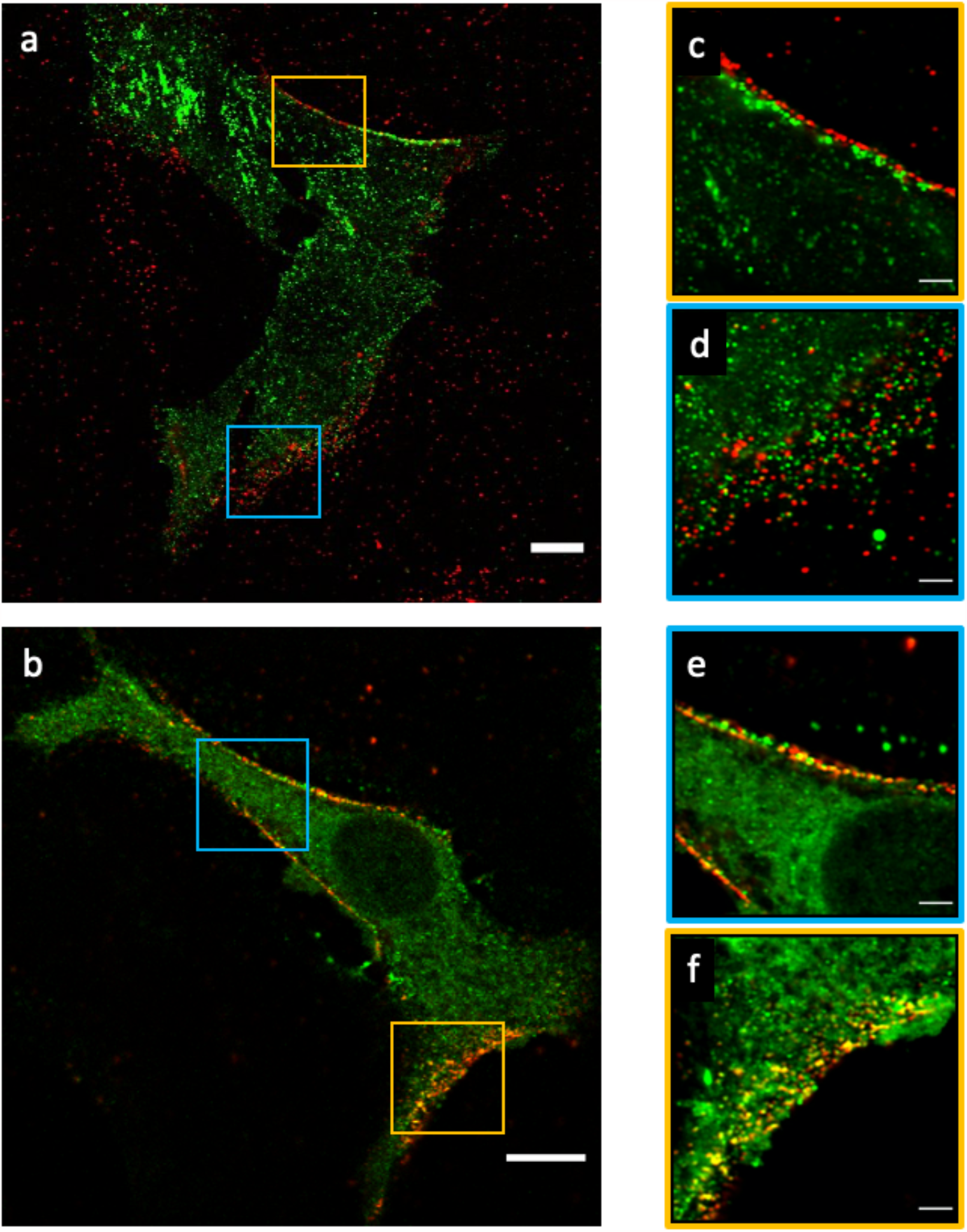
During the early entry phase virions on the membrane colocalize with clathrin but not with Caveolin-1. (a) Confocal images of virus (in red) with Caveolin-1 (in green); Green: Caveolin-1, red: S protein, scale bar: 10 µm. (b) Same as in (a) but Caveolin-1 is replaced by clathrin. (c-f) Airyscan images of regions in (a) or (b) enclosed in cyan and orange squares. Scale bar: 10 µm (a,b), 2 µm (c-f).

Perfect stoichiometric correlation (R=1) can never be achieved, owing to incomplete labeling, fluorescence background, and slight spatial mismatch of colors due to residual chromatic aberration. Accordingly, a positive control made of green/far-red doubly immunostained ACE2 receptor set the maximum achievable R to 0.69±0.01. With this reference, we found a medium/strong functional association of S with clathrin, but a poor or negligible association with caveolin-1 (Table 1).

Nanoscopy by dSTORM-TIRF at the basal membrane level demonstrated that single virions fully overlap with clathrin clusters (Figure 8). This supports clathrin-mediated endocytosis of the full virus, which was questioned by recent electron microscopy results in Vero E6 showing some clathrin pits at 50-100 nm from membrane-attached virions likely to endocytose released viral material^39^. A further support to clathrin-mediated late entry of SARS-CoV-2 was provided by τ-STED measurements, which addressed the apical submembrane level where most colocalized signal was visible (Figure 9). τ-STED images clearly showed single virions embedded into larger clathrin vesicles (170±90 nm) that, albeit not resolved into the structural triskelion, can safely be attributed to clathrin pits.

**Figure 8.**
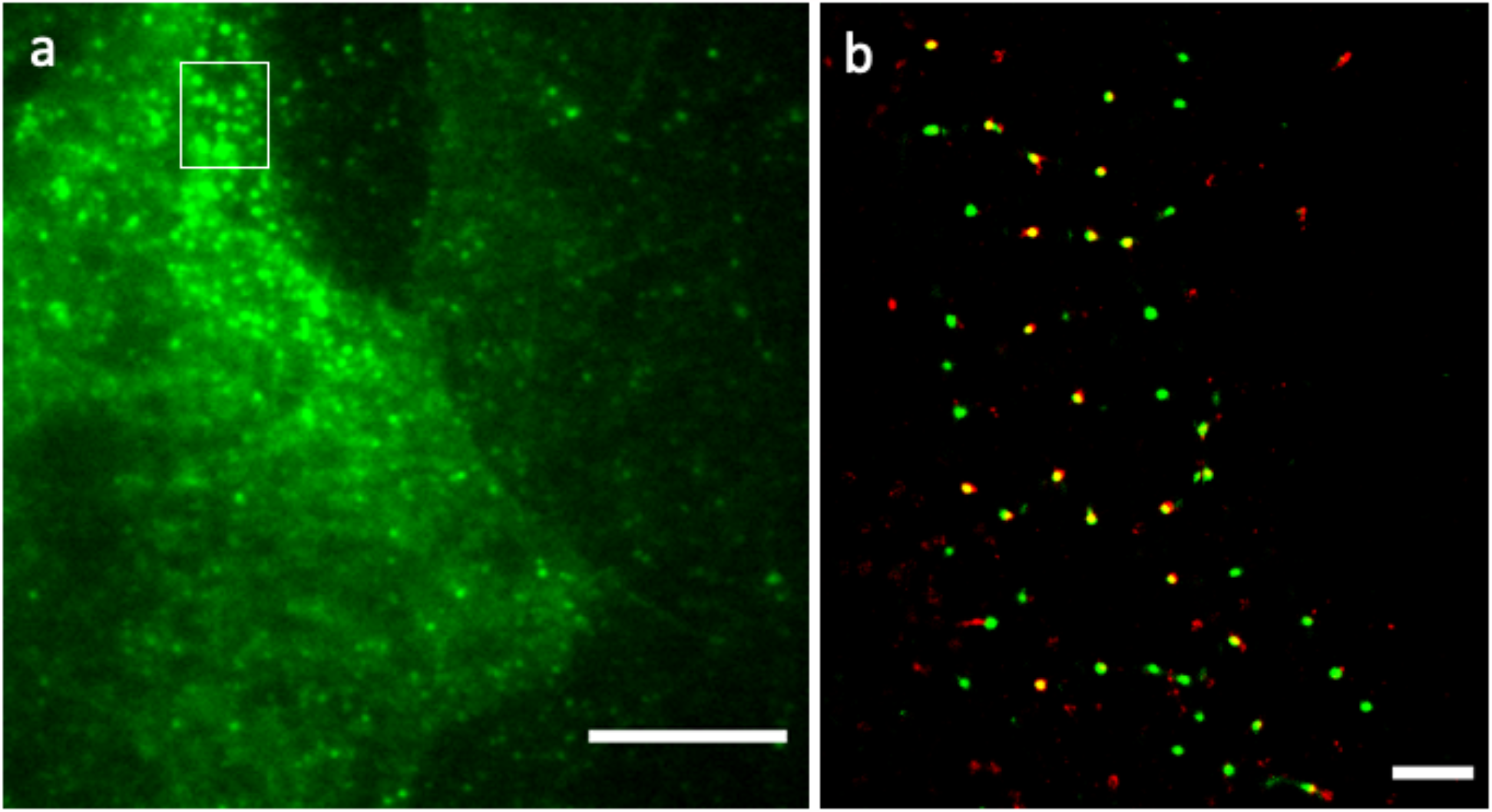
Virions are embedded into clathrin-coated pits. (a) TIRF image of Vero E6 cell after 3h post-incubation at 4 C (B.1.177 infection). (b) Two-color map of dSTORM-TIRF localization density of clathrin (green) and S (red): colocalized particles appear yellow; pixel size: 10 nm, average localization precision (±SD): 32±10 nm (both channels). Green: clathrin, red: S protein. Scale bar: 10 µm (a), 1 µm (b).

**Figure 9.**
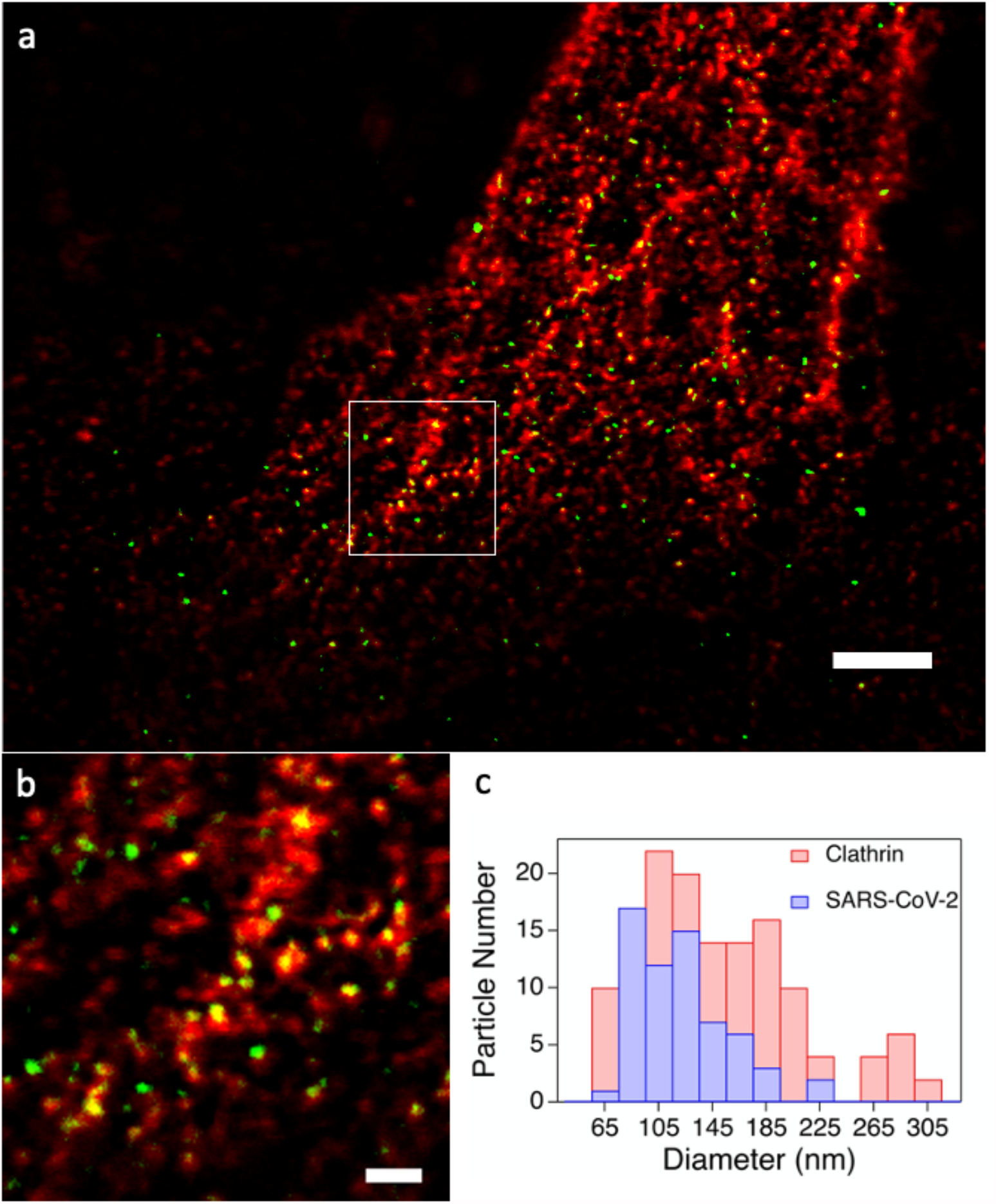
Virions are mostly embedded into submembrane clathrin-coated pits. (a) τ-STED image of clathrin and B.1.177 viruses attached to cell membranes; Green: S protein, red: clathrin. (b) Zoom of the region enclosed in the white square in (a). (c) Histogram of diameters (D) of clathrin (red) and viral (blue) particles found in (b): <D>=170+-90 nm (clathrin), <D>=120+-36 nm (virions). Scale bar: 2 µm (a), 200 nm (b).

The clathrin-mediated endocytosis of SARS-CoV-2 is at odds with the controversial hypothesis that ACE2 resides in caveolin-enriched raft regions of the cell membrane in several cell lines, including Vero E6^46, 47^. Accordingly, we set out to investigate the localization and functionality of ACE2 in the Vero E6 membrane by our microscopy toolbox. Confocal and TIRF imaging confirmed that ACE2 shows a prevalent membrane localization (Figure 10a), with some minor cytoplasmic staining. The functional receptor activity of membrane ACE2 towards SARS-CoV-2 was corroborated by the large colocalization with recombinant RBD of the S protein (Figure 10b, Table 1). Also, we found a significant degree of colocalization of ACE2 with CD71, the transferrin receptor (Figure 10b, Table 1). CD71 is known as a marker of the non-raft regions of the cell membrane^48^ and its clathrin-mediated endocytosis upon stimulation is well documented^49^. Conversely, Airyscan images highlighted negligible ACE2 colocalization with caveolin-1 (Figure 10c, Table 1).

**Figure 10.**
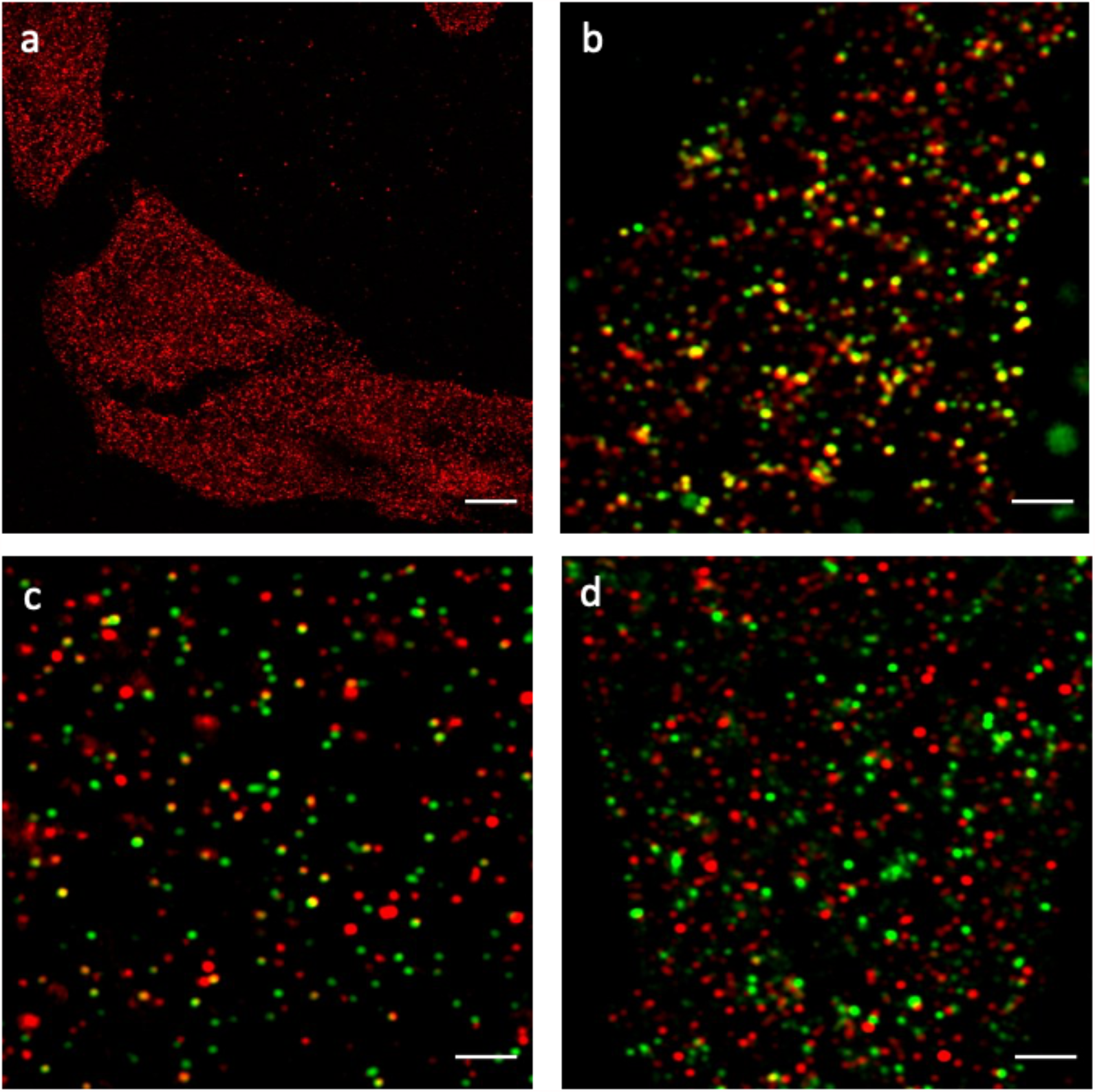
ACE2 and its molecular partners onto the membrane of Vero E6. (a) Membrane distribution of ACE2. (b) Airyscan colocalization image of ACE2 and SARS-CoV-2 RBD. (c) Airyscan colocalization image of ACE2 and Caveolin-1. (d) Airyscan colocalization image of ACE2 (red) and CD71 (green). Red: ACE2, green: RBD, Caveolin-1, CD71. Scale bar: 10 µm (a), 2 µm (b-d).

We can conclude that ACE2 localizes poorly in caveolin-1-enriched membrane regions in Vero E6 cells, in agreement with the absence of caveolar endocytosis of SARS-CoV-2. Additionally, the observed colocalization with CD71 supports the presence of ACE2 in non-raft regions of the cell membrane, wherefrom it may activate clathrin-mediated endocytosis upon contact with the S protein of SARS-CoV-2.

## DISCUSSION

The ongoing COVID-19 pandemic makes imperative the full understanding of virus-host interactions. In this context, the pivotal role of the surface Spike (S) protein was early recognized since ir mediates both the docking with the host cell receptor (ACE2) and the fusion process. The subtle interplay of S with the ACE2 receptor, its ability to hijack the cell endocytic machinery, and its intrinsic tunable fusogenic properties are directly related to the viral tropism. The SARS-CoV-2 S is the antigen encoded by available vaccines^1^. Also, the S glycoprotein represents the main target of therapeutic approaches aimed at neutralizing virus infectivity. S appears also key to viral adaptation to humans under selective pressure, and its sequence variability has already enabled the emergence of dominant viral variants harboring mutations such as the D614G present in lineage B.1.177 and, more recently, the Variants of Concern Alpha (B.1.1.7 lineage).

In spite of the accumulated knowledge, the thorough elucidation of unclear checkpoints of S-mediated entry requires a research approach focused more on single-virus interaction with the host cell^32^. In this perspective, for the first time in virus-cell studies we leveraged the combination of conventional microscopy (confocal, TIRF) with super-resolution microscopy techniques (Airyscan, STED, SMLM) whose common ability to image biological samples at <200 nm scale is properly tailored to the size of SARS-CoV-2. This toolbox enables the imaging of single-virus interactions with cells at different spatial scales, affording both functional and structural details. Of note, we applied our spatial multiscale imaging toolbox to the real virus, because virus models may not fully recapitulate the complex arrangement of S protein on virus envelope and its interaction with the target cells^50^.

The entry kinetics of B.1.177 and B.1.1.7 viral isolates was the first aspect evaluated in this study. While B.1.177 has been dominant in several European countries from summer 2020, the rapid spread of B.1.1.7 in the last months suggests a transmission advantage likely conferred by its genome changes. Several effects can contribute to improved transmissibility, including more rapid viral replication, faster entry, and the ability to escape the innate immunity of the host ^51^. Although present data are somewhat conflicting^52, 53^, a recent report hinted at the ability of P681H mutation of B.1.1.7 to influence cleavability at the furin S1/S2 site (Scheme 1) in TMPRSS2 expressing cells, thereby modulating the cell entry of the virus^54^. Yet the (verified) absence of TMPRSS2 on the surface of Vero E6 cells avoids the kinetic interference of protease-mediated priming and activation of S at the cell plasma membrane. Our findings clearly show that: 1) no major morphological differences exist between B.1.177 and B.1.1.7 viruses, 2) internalization of B.1.1.7 is much faster than B.1.177, 3) both strain gain access to Vero E6 cells through the “late pathway” i.e. via endosomal internalization and cathepsin-mediated cleavage and activation of S protein. Notably, the endosomal activity of cathepsins (particularly cathepsins B/L) occurs at proteolytic sites that are distinguishable and downstream of the furin cleavage site (e.g. ^694^AYTM^697^, see Scheme 1b)^55^, suggesting a minor role -if any-of the P681H mutation in accelerating B.1.1.7 access to cells. In this context, we are tempted to attribute the faster kinetic phenotype of B.1.1.7 to the N501Y replacement, which is thought to boost the binding interaction with the ACE2 receptor on the cell membrane^26, 27^ and could thereby trigger more rapid activation of the endosomal machinery.

The second question we addressed is the actual roles of clathrin and caveolin-1 in the late pathway of SARS-CoV-2. To our knowledge, no direct visualization of the protein mediator of endosomal entry followed by SARS-CoV-2 has been reported yet. Structural similarities with other CoVs such as NL63^56^ and SARS-CoV^57^ allegedly point out clathrin as the likeliest endocytic mediator of SARS-CoV-2 entry. Indeed, clathrin-mediated internalization of the naked S protein and S-pseudotyped lentivirus has been recently demonstrated^58^. Nonetheless, previous studies on SARS-CoV suggested an endocytic mechanism mediated by neither clathrin nor caveolin-1^59^. Our results clearly demonstrated the role of clathrin vesicles as major carriers of the virus from the surface to the early endosome. Conversely, caveolin-1 seems not to participate significantly in virus entry. This finding is in excellent agreement with the large exclusion of the ACE2 receptor from the caveolin-enriched raft domains of the cell membrane. The latter results are particularly interesting, as some authors alleged the role of lipid rafts and caveolin-1 in SARS-CoV entry^47, 60^. Yet, exclusion of ACE2 from these membrane regions was reported by other authors^61, 62^ and a major role of caveolin-1 in ACE2 distribution and virus entry is not easy to reconcile with the popular hypothesis of a clathrin-mediated first step of late entry in CoVs.

## CONCLUSIONS

A fluorescence microscopy imaging toolbox, which harnesses both conventional and super-resolution fluorescence microscopy and easily matches the spatial scale of single virus-cell checkpoints, was developed to tackle some major issues related to the entry of the full SARS-CoV-2 virus by a late pathway in Vero E6 cells. The B.1.1.7 variant that emerged in several countries showing a clear transmission advantage, as compared to its ancestor B.1.177 lineage that -mostly because D614G mutation in the S protein-became one of the dominant variants worldwide in 2020. Our results clearly indicate that B.1.1.7 outcompetes B.1.177 in terms of a much faster kinetics of entry in Vero E6 cells. This crucial property, perhaps one among the key factors of the evolutionary success of B.1.1.7, is likely unrelated to the P681H mutation. In fact, the latter might affect the furin cleavage site, but virus entry in Vero E6 is not mediated by TMPRSS2-fusion step at the cell membrane. Indeed, we demonstrated that both viral strains are internalized only by the “late pathway”, i.e. through the endosomal route and cathepsin-mediated activation of the S protein, which occurs at cleavage sites downstream of the furin site^55^. The faster cell entry of B.1.1.7 could be related to the stronger interactions of its S protein with the ACE2 receptor due to the N501Y substitution compared to B.1.177, the latter being representative of the original D614G clades which dominated in Europe from summer until late 2020.

A second result of our approach was the clear and direct demonstration of clathrin as the main mediator of endocytosis in the late pathway, a mechanism that had been previously suggested only by analogy with other CoVs and from experiments with pseudotyped virus models. Notably, our data also reconciled clathrin-mediated endocytosis with ACE2 localization on membranes, showing that the receptor does not colocalize with caveolar-enriched regions, as previously reported.

In conclusion, we believe that our fluorescence microscopy imaging toolbox represents a fertile strategy to address urgent questions on virus-cell checkpoints at the single virus level, while avoiding the conflicting results sometimes obtained using models unable to recapitulate the desired viral phenotypes.

## Author Contributions

BS, FB, AD, MP, RB designed the study; BS, PQ, NC, PB, GS, RB planned and performed research; CDP, NC, NM, EC, PGS, VC, GL, EP performed research; BS, PQ, CDP, NC, NM, EC, VC, EP, MC, PB, GS, RB analyzed data; BS, PQ, CDP, NC, GF, ML, MC, FB, AD, MP, RZ, PB, GS, RB wrote and revised the paper.

## Funding

This research was supported by MIUR, Progetto di Ricerca di Interesse Nazionale, (bando PRIN 2017, Project n. 2017KM79NN), Regione Lombardia-Fondazione CARIPLO, bando POR-FERS 2014-2020 (grant PAN-ANTICOVID-19). All funding sources had no involvement in study design, collection, analysis, interpretation of data, writing the report; and decision to submit the article for publication.

## Acknowledgments

Dr. Pasqualantonio Pingue (NEST, Scuola Normale Superiore) and Dr. Michele Oneto (IIT Nanophysics) are gratefully acknowledged for technical assistance and support.

## MATERIALS AND METHODS

### Cell lines and culture

African green monkey kidney cells (Vero E6) were obtained from ATCC (CRL-1586). Vero E6 were cultured in DMEM high glucose medium supplemented with heat-inactivated 10% fetal bovine serum (FBS) (Sigma-Aldrich, Milan, Italy), 2 mM L-glutamine, 10 U/ml penicillin, and 10 mg/ml streptomycin (Sigma-Aldrich, Milan, Italy), at 37°C in the presence of 5% CO_2_.

### Virus isolation and amplification

Clinical isolates of SARS-CoV-2, B.1.177 and B.1.1.7 (hCoV-19/Italy/LOM-UniSR10/2021, GISAID Accession ID: EPI_ISL_2544194 and hCoV-19/Italy/LOM-UniSR7/2021, GISAID Accession ID: EPI_ISL_1924880) were isolated on Vero E6 cells from nasopharyngeal swabs from COVID-19 patients in BSL-3 facility at Laboratory of Microbiology and Virology, Vita-Salute San Raffaele University. All the procedures of infection were performed in a BSL3 facility. An aliquot (0.8 mL) of the transport medium of the nasopharyngeal swab (COPAN’s kit UTM® universal viral transport medium—COPAN) was mixed with an equal volume of DMEM without FBS and supplemented with a double concentration of P/S and Amphotericin B. The mixture was added to 80% confluent Vero E6 cells monolayer seeded into a 25 cm^2^ tissue culture flask. After 6 h adsorption at 37°C, 1 mL of DMEM supplemented with 2% FBS and Amphotericin B were added. Twenty-four hours post-infection (hpi), 3 mL of DMEM supplemented with 2% FBS and Amphotericin B were added after a PBS wash. Live images were acquired (Olympus CKX41 inverted phase-contrast microscopy) daily for evidence of cytopathic effects (CPE), and aliquots were collected for viral RNA extraction and In-house one-step real-time RT-PCR assay. Five days post-infection (dpi) cells and supernatant were collected, aliquoted, and stored at -80°C (P1). For secondary (P2) virus stock, Vero E6 cells seeded into 25 cm^2^ tissue culture flasks were infected with 0.5 mL of P1 stored aliquot, and infected cells and supernatant were collected 48 hpi and stored at -80°C. For tertiary (P3) virus stock, Vero E6 cells seeded into 25 cm^2^ tissue culture flasks were infected with 0.2 mL of P2 stored aliquot and prepared as above described.

### Virus titration

P3 virus stocks were titrated using both Plaque Reduction Assay (PRA, PFU/mL) and Endpoint Dilutions Assay (EDA, TCID_50_ /mL). For PRA, confluent monolayers of Vero E6 cells seeded in 24-well plates were infected with 10-fold-dilutions of virus stock. After 1 h of adsorption at 37°C, the cell-free virus was removed, and cells washed with PBS. Cells were then incubated for 46 h in DMEM containing 2% FBS and 0.5% agarose. After fixing and staining the cell monolayers, viral plaques were counted. For EDA, Vero E6 cells (3·10^5^ cells/mL) were seeded into 96-well plates and infected with base 10 dilutions of virus stock. After 1 h of adsorption at 37°C, the cell-free virus was removed, and complete medium was added to cells after a PBS wash. After 72 h, cells were observed to evaluate CPE. TCID_50_/mL was calculated according to the Reed–Muench method.

### Kinetic study of virus growth in cells

Vero E6 cells (3·10^5^ cells/mL) were seeded into 96-well plates and cultured for 1 day at 37°C. Then, two experimental settings, characterized by a long exposure or a short exposure to the virus, were performed in triplicate. In the first experiment, the cells were infected for 6, 24, and 48 h with B.1.177 or B.1.1.7 at 0.001 MOI, and cell supernatants were collected at the different time points. In the short exposure experiment, the cells were washed three times with PBS after 1 h of virus adsorption (B.1.177 or B.1.1.7 at 0.001 MOI), and cells were incubated for 1, 3, 6, 12, 18, 24, 48, and 72 hpi. The cell supernatants were collected at the different time points as well. Viral genome from the supernatants of both experimental settings was extracted and analyzed by real-time RT-PCR.

### Inhibition of SARS-CoV2 infection by E64d

Vero E6 cells (3·10^5^ cells/mL) were seeded into 96-well plates and cultured for 1 day at 37°C. Subsequently, cells were incubated 1h before infection with 10 µM E64d, a Cathepsin L inhibitor, (Sigma Aldrich, Milan, Italy). Then, cells were infected for 1h with B.1.177 or B.1.1.7 (0.001 MOI) virus. After virus adsorption, the cells were washed three times with PBS and incubated for 48 h with 10 µM E64d. Cells infected without E64d treatment were added as well as experimental control. The experiment was performed in triplicate, and the cell supernatants were collected for viral genome extraction and subsequent analysis by RT-PCR.

### Real-time RT-PCR of SARS-CoV-2

SARS-CoV-2 RNA relative amounts detected for each experimental condition as a cycle threshold (Ct) value were compared, with a mean Ct value determined for the positive infection control. The viral RNA was purified from 100 μL of all cell-free culture supernatant, using the QIAamp Viral RNA Mini Kit (Qiagen). The purified RNA was then used to perform the synthesis of first-strand complementary DNA, using the SuperScript First-Strand Synthesis System for RT-PCR (Thermo Fisher Scientific).

Real-time PCR, using the SYBR Green dye-based PCR amplification and detection method, was performed to detect the complementary DNA. We used the SYBR Green PCR Master Mix (Thermo Fisher Scientific), with the forward primer N2F (TTA CAA ACA TTG GCC GCA AA), the reverse primer N2R (GCG CGA CAT TCC GAA GAA). The PCR conditions were 95°C for 2 minutes, 45 cycles of 95°C for 20 seconds, annealing at 55°C for 20 seconds and elongation at 72°C for 30 seconds, followed by a final elongation at 72°C for 10 minutes. RT-PCR was performed using the ABI-PRISM 7900HT Fast Real-Time instrument (Applied Biosystems) and optical-grade 96-well plates. Samples were run in duplicate, with a total volume of 20 μL.

### Analysis of TMPRSS2 expression by real-time RT-PCR

Vero E6 cells were seeded in 6-well plates at 2·10^5^ cells/well and cultured for 1 day at 37°C. Three wells were washed once with PBS, after that 1ml TRIZOL (Thermo Fisher Scientific) was added. Samples were then transferred in 4 ml tubes and processed for total RNA extraction with RNA micro kit (ZYMO REAGENT) according to company protocol. RNA quantity and integrity were assessed with Qubit 4.0 fluorometer (Thermo Fisher Scientific) using Qubit RNA BR kit (Thermo Fisher) and Qubit RNA IQAssay kit respectively. 500 ng of each sample were reverse transcribed with iScript gDNA clear cDNA Synthesis kit (BioRad) according to kit protocol and 10 ng of cDNA were analyzed for TMPRSS2 expression by real-time RT-PCR on a CFX Connect Real Time System using SsoAdvancedSybrGreen Supermix (BioRad). The amplification protocol was 2 min at 95°C, 40 two-step cycles of 10 s at 95°C and 30 s at 60°C, final ramping from 65°C to 95°C with 0.5°C increments at 5 sec/step, for amplicon melting profile. The cDNA of Caco-2 and HepG2 cells was used as a positive control of TMPRSS2 amplification. All values were normalized by the housekeeping gene RPL13A. All samples were run in duplicate.

### Cell infection (unsynchronized) for immunofluorescence study of virus entry

10^5^ Vero E6 cells were seeded in 35 mm glass bottom dishes (Willco, Amsterdam) with 2 ml of culture medium and cultured for 1 day at 37°C. Subsequently, the medium was removed, and the cells were inoculated for 1, 3, 6, or 72h with B.1.177 or B.1.1.7 at 0.001 MOI while keeping the temperature at 37°C. At the end of incubation time, the medium was removed, the cells were washed 3 times with 500 µl of PBS and then were fixed and permeabilized with ice-cold 100% methanol (Sigma Aldrich) for 15 minutes at -20°C. After methanol was removed, cells were rinsed again three times in PBS for 5 minutes each.

### Cell infection (synchronized) for immunofluorescence study of virus endocytosis

10^5^ Vero E6 cells were seeded in 35 mm glass bottom dishes (Willco, Amsterdam) with 2 ml of culture medium and cultured for 1 day at 37°C. Cells were then pre-chilled by incubation at 4°C for 30 min. Subsequently, the medium was removed, and cells were infected at MOI 2 for 3 h with B.1.177 keeping the temperature at 4°C. At the end of incubation, the viral inoculum was removed, cells were gently washed with ice-cold PBS, and cell culture medium was added. Cells were incubated at 37°C and 5% CO_2_ for 3 h. Next, the medium was removed, cells were washed 3 times with 500 µl of PBS, and finally fixed and permeabilized with ice-cold 100% methanol for 15 minutes at -20°C. After methanol was removed, cells were rinsed again three times in PBS for 5 minutes each.

### Primary antibodies for immunofluorescence studies

- anti-S IgG rabbit monoclonal antibody (40592-V05H, Sino Biological), dilution: 1:200
- anti-N IgG rabbit monoclonal antibody (40143-R019, Sino Biological), dilution: 1:200
- anti-ACE2 IgG rabbit monoclonal antibody (ab15348, AbCam), dilution: 1:200
- anti-clathrin IgG mouse monoclonal antibody (sc-12734, SantaCruz), dilution: 1:200
- anti-caveolin-1 IgG mouse monoclonal antibody (sc-70516, SantaCruz), dilution: 1:200
- anti-CD71 IgG mouse monoclonal antibody (sc-65882, SantaCruz), dilution: 1:100
- anti-α-tubulin IgG mouse monoclonal antibody (T5168, Merck), dilution: 1:1000

### Secondary antibodies for immunofluorescence studies and combinations

- donkey anti-rabbit IgG Alexa488-labeled monoclonal antibody (a21206, ThermoFisher), dilution: 1:500 (confocal, airyscan, and dSTORM-TIRF experiments)
- donkey anti-rabbit IgG Alexa647-labeled monoclonal antibody (a31573, ThermoFisher), dilution: 1:500 (confocal, airyscan, and dSTORM-TIRF experiments)
- donkey anti-mouse IgG Alexa488-labeled monoclonal antibody (a21202, ThermoFisher), dilution: 1:500 (confocal, airyscan, and dSTORM-TIRF experiments)
- donkey anti-mouse IgG Alexa647-labeled monoclonal antibody (a31571, ThermoFisher), dilution: 1:500 (confocal, airyscan, and dSTORM-TIRF experiments)
- goat anti-rabbit IgG Atto647N-labeled monoclonal antibody (40839, Merck), dilution: 1:200 (τ-STED experiments)
- goat anti-mouse IgG Atto594-labeled monoclonal antibody (76085, Merck), dilution: 1:200 (τ-STED experiments)

### Immunostaining of infected cells

Methanol-fixed infected cells and a methanol-fixed negative control were incubated overnight at 4 C° with 150 μl of a solution of anti-S IgG or anti-N IgG in PBS + 3% BSA (Sigma-Aldrich, Milan, Italy). For colocalization experiments, the incubation solution was supplemented with anti-clathrin IgG or anti-caveolin-1 IgG. In one experiment on cells at 72 hpi, the incubation solution was supplemented with anti-tubulin IgG. After rinsing four times with PBS + 0.5% BSA (PBB), infected cells and negative controls were incubated for 1h with a solution of 1-2 fluorescently-labeled secondary antibody/ies in PBB (see secondary antibody section in Materials and Methods) and then rinsed four times with PBB and three times with PBS. When required, cell nuclei were stained by exposure for 5 min to 1 mg/100 ml Hoechst 33342 (ThermoFisher) in water.

### Immunostaining of non-infected cells

10^5^ Vero-E6 cells were seeded in 35 mm glass bottom dishes (Willco) with 2 ml of culture medium and cultured for 1 day at 37°C. Then, cells were fixed with PFA 2% in PBS for 15 min, rinsed three times with PBS, permeabilized for 15 min with Triton-X 100 (Sigma) 0.1% in PBS for 15 min, and rinsed four times with PBS + 0.5% BSA (PBB). Fixed cells were incubated overnight at 4 C° with 200 μl of a solution of anti-ACE2 IgG, and either Spike RBD-mFC Recombinant Protein (40592-V05H-100, SinoBiological), or anti-caveolin-1 IgG, or anti-CD71 IgG. After rinsing four times with PBB, immunolabeled cells and negative controls were incubated for 1h with a solution of 2 fluorescent-labeled secondary antibodies in PBB (see secondary antibody section in Materials and Methods) and then rinsed four times with PBB and three times with PBS. When required, cell nuclei were stained by exposure for 5 min to 1 mg/100 ml Hoechst 33342 (Thermo Fisher Scientific) in water.

### Confocal and Airyscan microscopy

Fluorescence was measured by a confocal Zeiss LSM 880 with Airyscan (Carl Zeiss, Jena, Germany), supplied with GaAsP detectors (Gallium:Arsenide:Phosphide). Samples were viewed with a 63x Apochromat NA=1.4 oil-immersion objective. We adopted 0.9x zoom for imaging multiple cells (1 pixel = 220 nm), and 2-6x zoom for imaging single cells; Airyscan imaging was carried out at zoom >3. The pinhole size was set to 44 mm, which corresponds to 1 airy unit (AU) for the green acquisition channel. Pixel dwell time was adjusted to 1.52 us and 512×512 pixel or 1024×1024 images were collected. In confocal mode, we carried out concomitant acquisition for all channels line by line with line-average set to 4. In airyscan mode, we carried out a sequential acquisition for all channels with frame-average set to 4. The acquisition channels were set as follows:

- Blue (Hoechst 33342): λ_ex_ =405 λ_em_ = 420-500 nm
- Green (Alexa488): λ_ex_ =488, λ_em_ = 500-560 nm
- Far-red (Alexa647): λ_ex_ =640, λ_em_ = 650-700 nm

Images were visualized and processed by the open-source software Fiji (NIH, Bethesda). Colocalization of the green and far-red images was quantified by Pearson’s coefficient *R* according to the method by Costes et al. ^63^ by the *colocalization threshold* and *colocalization test* routines of Fiji.

### Single Molecule Localization by dSTORM-TIRF

A commercial N-STORM TIRF microscope (Nikon Instruments), equipped with an oil immersion objective (CFI Apo TIRF 100×, NA 1.49, oil; Nikon) was used to acquire 40,000 frames at a 33 Hz frame rate using TIRF illumination. Excitation intensities were as follows: ∼ 0.5-1 KW/cm^2^ for the 647 nm readout (200 mW laser; MPB Communications), ∼ 0.1-0.2 KW/cm^2^ for the 488 nm readout (50 mW laser; Oxxius), and ∼ 35 W/cm^2^ for the 404 activation (100 mW laser; Coherent). For single color measurements, we set a repeating cycle of 1 activation frame at 404 nm / 3 readout frames at 647 nm or 488 nm. For double color measurements, we set a repeating cycle of 1 activation frame at 405 nm / 3 readout frames at 488 nm / 1 activation frame at 405 nm / 3 readout frames at 647. Image detection was performed with an EMCCD camera (Andor iXon DU-897; Andor Technologies) with EM gain activated and set to 300. We set full TIRF excitation of the sample by changing the objective back-aperture illumination through the acquisition software of the Microscope (NIS Elements AR 5.20.01, Nikon). The Perfect Focus System (Nikon) was used during the entire recording process. Fluorescence-emitted signal was spectrally selected by the four-color dichroic mirrors (ZET405/488/561/647; Chroma) and filtered by a quadribandpass filter (ZT405/488/561/647; Chroma).

For imaging conditions, STORM imaging buffer was used containing a glucose oxidase solution as an oxygen scavenging system. Imaging buffer was prepared as follows. 690 uL of 50mM Tris buffer (pH 8.0), containing 10mM of NaCl and 10% w/v of glucose, were mixed with 25 uL of DL lactate (60% w/w syrup in water, Sigma Aldrich, L1375-100ML) and 3.5 uL of COT (200 mM in DMSO). The solution was stored at 4 C and filtered (220 nm) before use. Immediately prior to the use, the solution was mixed with 3.5 uL of GLOX solution, 3.5 uL of 2-mercaptoethanol, 25 uL of Cysteamine (1M in H_2_O, Sigma Aldrich, 30070-10G), and 45 uL of Oxyrase. The resulting solution was added to the petri dish, which was sealed with aluminum tape. GLOX solution was composed of glucose oxidase (14 mg) and Catalase (50 uL, 17 mg/mL) dissolved in buffer A (200 uL). GLOX solution was stored at 4 C for a maximum of 14 days.

### Single molecule localization analysis

Acquired dSTORM stacks were processed by Thunderstorm, a Fiji plugin for PALM and STORM data analysis ^64^. At first, we set the properties of acquisition by the “Camera setup” menu: pixel size = 158.7 nm, Photoelectrons per A/D count: 2.5, Base level: 100 counts, EM gain: 300. Then, we carried out the localization algorithm (“Run analysis”), setting the following parameters: a) pre-filter: difference of averaging filters with 3 and 6 pixels as first and second kernel size, respectively; b) approximate localization of molecules by local maximum method with threshold 200 and 8-neighbourhood connectivity; c) sub-pixel localization by the Integrated Gaussian method, performing least-squares multi-fitting (threshold *p*=1E-6) with initial sigma 1.6 pixels and fitting radius 3 pixels, maximum 5 molecules for fitting region with limit intensity range 1-1000 photons. Eventually, we cleaned the obtained results from drift and those localizations not strictly lying on the focal plane by the following post-filtering algorithm: a) removal of first 500 frames; b) drift correction by correlation; c) merging reactivated molecules (max distance: 20 nm, max off frames: 1, limited frames per molecule); d) removal of localizations with: (intensity = 1000 AND sigma >180 nm AND uncertainty > 130 nm).

### Single molecule localization density maps and cluster analysis

Single molecule localization maps and cluster analysis were collected by the LocAlization Microscopy Analyzer software (LAMA), available for download at http://share.smb.uni-frankfurt.de/index.php/software-menue/lama. Before LAMA analysis, the localization list exported by Thunderstorm was converted to the Molecular Accuracy Localization Keep (MALK) format used by the LAMA by the localization converter routine of the LAMA software.

Single molecule localization density maps were obtained by the Visualization routine of LAMA and consisted of 2D histograms of the localization list obtained by using a pixel size of 10×10 nm and codified into a 0-255 color map. For dual color images, before visualization, the localization lists of both colors were spatially registered by the register cabinet of the LAMA by using localization lists of multicolor beads. This procedure is extensively described in the documentation file accompanying the LAMA software, which can be downloaded at https://share.smb.uni-frankfurt.de/index.php/component/jdownloads/download/4-lama-tutorial/8-lama-documentation. Hierarchical Cluster Analysis (HCA) was performed by the Density-Based Algorithm for Discovering Clusters in Large Spatial Databases with Noise (DBSCAN) preceded by Ordering Points To Identify the Clustering Structure (OPTICS) algorithm ^42^. OPTICS-DBSCAN was performed an 8×8 um Region of Interest (ROI) (3-5 for each cell) by setting the minimum cluster size to 5 localizations and the noise level to 10%.

### *Stimulated Emission Depletion Microscopy (*τ-STED)

Lifetime-tuning STED (τ-STED) measurements were performed using a Leica STELLARIS 8 Falcon τ-STED (Leica Microsystems, Mannheim, Germany) inverted confocal/STED microscope. Excitation was provided by a White Light Laser and selecting the following wavelengths by the acoustic-optical tunable filter (AOTF): 488 nm, 560 nm, and 638 nm. Detection has been performed by the embedded tunable spectrometer in the 500 - 550 nm, 570 - 630 nm, 660-750 nm ranges respectively, and three Power HyD detectors. The pinhole was set to 0.6–1 Airy size. Line scanning speed ranged from 10 to 1400 Hz in standard acquisition mode. In τ-STED mode, the 775 nm pulsed laser beam is superimposed at a typical power of 100 – 250 mW before the objective. Two-colors τ-STED has been performed sequentially by line for the red and far-red fluorophores. Green fluorophores are not affected by the depletion beam at 775nm.

### Graphics and statistics

Graphs were prepared using Prism 7 (GraphPad) and IgorPro8 (Wavemetrics) software. Data are shown as the mean +/- SEM. Statistical analysis was performed by Prism 7 (GraphPad). For comparisons amongst Cycle thresholds (Ct) the 2way ANOVA multiple comparisons with corrections were performed by using Prism 9 (GraphPad).

## Notes

### Competing Interest Statement

The authors have declared no competing interest.

