## Supplementary Information for "A spatial multi-scale fluorescence microscopy toolbox discloses entry checkpoints of SARS-CoV-2 variants in Vero E6 cells"

***Numerical simulation of S localization density under TIRF excitation***

Our system is based on the schematic arrangement of a viral particle in the evanescent field of the TIRF microscope (Scheme S1). In more details, the envelope of the viral particle is at a distance d from the glass-water interface (coverslip) where the evanescent field is maximum. The field decays mono-exponentially along the z direction with a spatial constant of 100 nm. Each S protein on the virus surface has 25 nm length and is inclined by 40 degrees with respect to the normal. We assume that the epitope recognized by the primary anti-S antibody (Ab1) is at distance ℓ from the point where S emerges from the envelope (Scheme 1S, inset: blue two-way arrow). For visualizing the differences, we set ℓ=12.5 (mid-S protein), 16.7 , and 20.8 nm. A secondary, fluorescent-labeled antibody is bound to Ab1: the exact position of the fluorescent labels with respect to the S stalk is unknown, but we may assume an average distance of 20 nm from S (Scheme 1S, inset: red two-way arrow) summing the 17.5 nm enlargement effects on microtubule imaging described by[1] and the half-thickness (2.5 nm) of the upper part of S protein itself (Scheme 1S, inset: black two-way arrow).

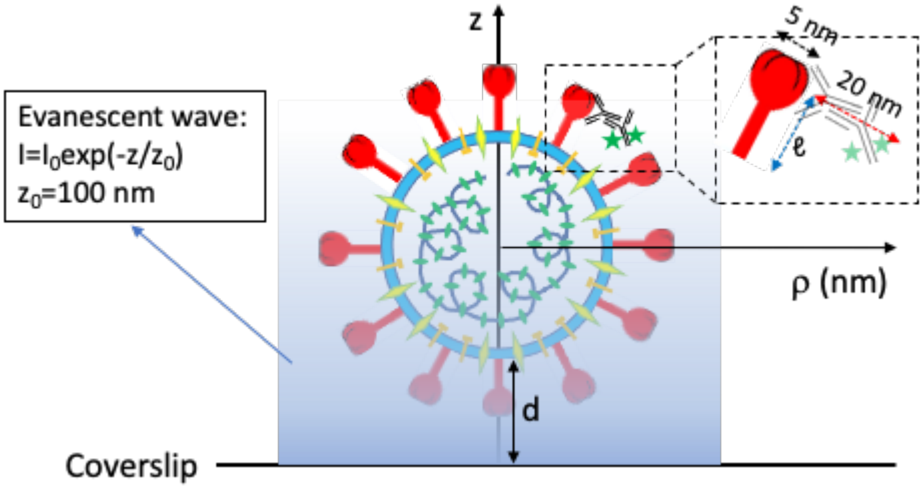


**Figure S1. Scheme of a viral particle, labeled on the S protein by a primary/secondary (fluorescent) antibody couple, excited by the TIRF evanescent field.** (a) Scheme of excitation: the viral particle is displaced by distance d from the coverslips and is flooded by an evanescent excitation wave that has cylindrical symmetry around the z-axis and decays monoexponentially along the z-axis with spatial constant 100 nm. The z-axis crosses the virus center, and the displacement from the virus center is measured by the radial coordinate ρ (nm). Inset: the details of S labeling by the couple primary-secondary antibody is shown. ℓ is the distance of the recognized epitope from the basis of S, and we assume that ℓ =12.5, 16.7, 20.8 nm, i.e. three different positions that cover the second-half of the S protein. The average distance of the fluorescent label from the epitope (supposed along the S axis) is 20 nm, adding half of the thickness (2.5 nm) of the upper part of S protein to the average enlargement of single molecule localizations in microtubules found by [1].

We calculated the position of the fluorophore in the space, i.e. of the single molecule to be localized, a Montecarlo simulation gathering all the described features. In more details, at first we designed a sphere of radius 45 nm centered in x=y=0, z=45 nm + d (with d=0, 10, 25 and 50 nm) simulating the viral envelope at variable distance from the coverslip. Then, in each iteration we set a random point on the sphere surface and from it we draw a line segment L defining an angle of 40 degrees with respect to the normal to the sphere. Subsequently, we move at a distance ℓ=12.5, 16.7, 20.8 nm along L. Eventually, we move further 20 nm in a random direction orthogonal to L. The final point P(x,y,z) represents one accessible position of the average fluorescent label bound to the secondary antibody. On account of the cylindrical symmetry of the system, the (x,y) coordinates can be replaced by the radial distance $\rho = \mathrm{sqrt}(x^2+y^2)$, yielding P(ρ,z). All points P with the same rho are accumulated, each one weighted by the probability of excitation $[\exp(z/z_0)]$ to obtain a distribution which is finally divided by 2πρ, to normalize along a radial section. This distribution corresponds to the localization map of single molecules theoretically achievable by STORM with infinite precision of localization. To obtain the distribution corresponding to the experimental localization map, we convolved the distribution with a gaussian function with sigma = (30 nm)/sqrt(2) (Figure S2). Significantly, the distribution is rather flat up to ρ = 48-58 nm, which represents the radius of the viral particle observable by our dSTORM-TIRF experiments.

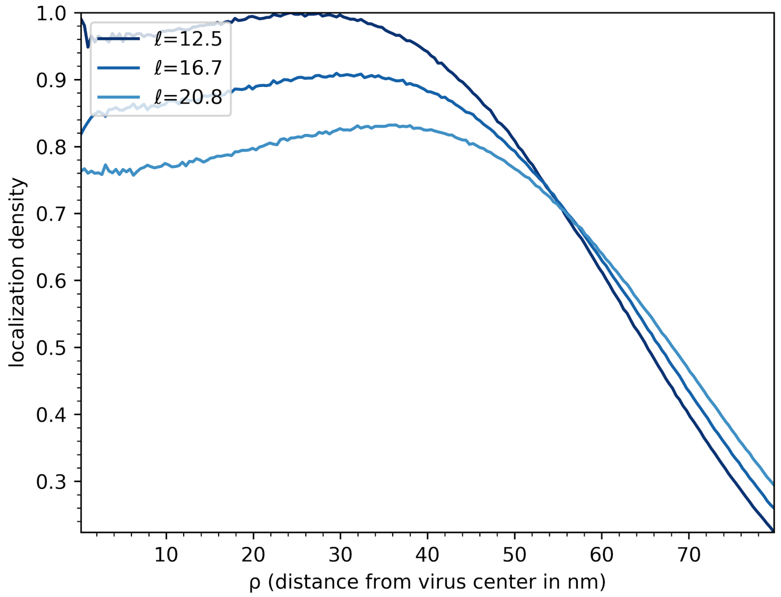


**Figure S2. Simulation of detectable S density by using a TIRF imaging system.** (a) Density of S protein localization along the radial coordinate ρ. The normalized distribution from Montecarlo is obtained from 100 million random points and the ρ value ranges from ρ = 0 to ρ = 80 nm divided into 200 equal bins.


1. Pleiner T, Bates M, Gorlich D. A toolbox of anti-mouse and anti-rabbit IgG secondary nanobodies. *J Cell Biol* **2018** 217, 1143. 10.1083/jcb.201709115.
